# Time-lagged Independent Component Analysis of Random Walks and Protein Dynamics

**DOI:** 10.1101/2021.03.18.435940

**Authors:** Steffen Schultze, Helmut Grubmüller

## Abstract

Time-lagged independent component analysis (tICA) is a widely used dimension reduction method for the analysis of molecular dynamics (MD) trajectories and has proven particularly useful for the construction of protein dynamics Markov models. It identifies those ‘slow’ collective degrees of freedom onto which the projections of a given trajectory show maximal autocorrelation for a given lag time. Here we ask how much information on the actual protein dynamics and, in particular, the free energy landscape that governs these dynamics the tICA-projections of MD-trajectories contain, as opposed to noise due to the inherently stochastic nature of each trajectory. To answer this question, we have analyzed the tICA-projections of high dimensional random walks using a combination of analytical and numerical methods. We find that the projections resemble cosine functions and strongly depend on the lag time, exhibiting strikingly complex behaviour. In particular, and contrary to previous studies of principal component projections, the projections change non-continuously with increasing lag time. The tICA-projections of selected 1 *μ*s protein trajectories and those of random walks are strikingly similar, particularly for larger proteins, suggesting that these trajectories contain only little information on the energy landscape that governs the actual protein dynamics. Further the tICA-projections of random walks show clusters very similar to those observed for the protein trajectories, suggesting that clusters in the tICA-projections of protein trajectories do not necessarily reflect local minima in the free energy landscape. We also conclude that, in addition to the previous finding that certain ensemble properties of non-converged protein trajectories resemble those of random walks, this is also true for their time correlations. Due to the higher complexity of the latter, this result also suggests tICA analyses as a more sensitive tool to test MD simulations for proper convergence.

## 1 Introduction

The atomistic dynamics of proteins, protein complexes, and other biomolecules is exceedingly complex, covering time scales from sub-picoseconds to up to hours [1, 2]. It is governed by a similarly complex high-dimensional free energy landscape or funnel [3], characterized by a hierarchy of free energy barriers [4], and has been widely studied computationally by molecular dynamics (MD) simulations [5]. With particle numbers ranging from several hundreds to hundreds of thousands or more [6, 7, 8, 9], the correspondingly high-dimensional configuration space of the system poses considerable challenges to a fundamental understanding of biomolecular function, e.g., of the conformational motions of these biological ‘nano-machines’ [10, 11], protein folding [12], or specific binding.

Several attempts to reduce the dimensionality of the dynamics have addressed this issue. Most notable approaches are principal component analysis (PCA) to extract the essential dynamics [13] of the protein that contributes most to the atomic fluctuations, and time-lagged independent component analysis (tICA), which identifies those collective degrees of freedom that exhibit the strongest time-correlations for a given lag-time [14, 15]. Both dimension reduction techniques can yield information on the conformational dynamics of a protein, i.e., how the protein moves through several conformational substates, which can be defined as metastable conformations characterized by local free energy minima [16].

This property also renders these dimension reduction techniques highly useful as a preprocessing step to describing the conformational dynamics of macromolecules in terms of a discrete Markov process [17, 18, 19]. Currently tICA is most widely used, and it is preferred over PCA for this purpose [20] because it additionally uses time information of the input trajectory.

In this context, both PCA and tICA rely on MD trajectories as input, which raises the question how much of these analyses is determined by actual information on the protein dynamics, as opposed to noise due to the inherently stochastic nature of each trajectory, and, importantly, how these two can be quantified.

For PCA, this question has been answered by analysis of the principal components of a high-dimensional random walk in a flat energy landscape [21, 22]. Unexpectedly, these turned out to approximate cosine functions, thus providing a very powerful criterion for the convergence of MD trajectories: The more an MD trajectory resembles a cosine, quantified by the cosine content [21], the more it resembles a random walk, and the less information it contains on the actual protein dynamics or the underlying free energy landscape.

These analyses [21, 22] have also suggested that clusters observed in low-dimensional PCA projections do not necessarily imply the existence of conformational substates and, instead, may also be a stochastic and/or projection artefact. Particularly the latter finding is highly relevant for the use of PCA for the construction of Markov models [19], which thus may also in part reflect the randomness of one or several trajectories. Note that this holds also true — albeit probably to a lesser extent — for the construction of Markov models from several or many trajectories, as these have to be spawned from a seeding trajectory or from starting structures generated from other advanced sampling methods [16, 23, 24, 25].

For tICA, no such analysis is available, but inspection of several examples suggests that similar effects may also be at work [26, 27]. To address this issue, here we will therefore analyze the tICA-projections of high dimensional random walks, and subsequently compare them to tICA-projections of selected protein trajectories. In particular, we will semi-analytically derive an expression for random walk tICA-projections, which will prove analogous to the PCA cosine functions and thus can also serve as a criterion for convergence as well as for the quality of derived Markov models. Unexpectedly, and contrary to the regular behaviour of random walk PCA projections, tICA-projections turn out to display much more complex behaviour. In particular, we observed critical lag times at which the random walk projections change drastically and — for high dimensions — even discontinuously. The resulting much richer and more intricate structure of random walk projections renders the proper interpretation of tICA-projections of protein dynamics trajectories particularly challenging, and has profound implications for the proper constructions of Markov models.

## 2 Theoretical Analysis and Methods

### 2.1 Definition of tICA

To establish notation, we briefly summarize the basic principle of tICA; for a more comprehensive treatment with particular focus on molecular dynamics applications, see Ref. [28].

Consider a *d*-dimensional trajectory **x**(*t*) = (*x*_1_(*t*), … , *x*_*d*_(*t*))^*T*^ ∈ ℝ^*d*^ with Cartesian coordinates *x*_1_, … , *x*_*d*_, which for compact notation we assume to be mean-free, that is, the time average ⟨**x**(*t*)⟩_*t*_ is zero. TICA determines those ‘slowest’ independent collective degrees of freedom **v**_*k*_ ∈ ℝ^*d*^, *k* = 1, … , *d*, onto which the projections *y*_*k*_(*t*) = **v**_*k*_ · **x**(*t*) have the largest time-autocorrelation

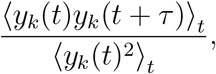

where *τ* is a chosen lag time. Equivalently, using the time-lagged covariance matrix

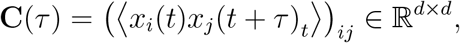

each degree of freedom **v**_*k*_ maximizes

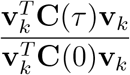

under the constraint that it is orthogonal to all previous degrees of freedom. Hence, the **v**_*k*_ are the solutions of the generalized eigenvalue problem

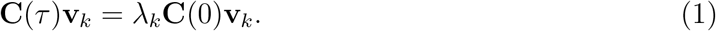

We will use the term ‘tICA-eigenvector’ for the v_*k*_ and ‘tICA-projection’ for the projections *y*_*k*_ onto the tICA-eigenvectors. In the literature, the term ‘tICA-component’ is often used, but it is somewhat ambiguous and we will therefore avoid it.

For an infinite trajectory of a time-reversible system the matrices in this eigenvalue problem are symmetric. However, for the finite trajectories considered here, with time steps *t* = 1, … , *n*, the matrix **C**(*τ*) is usually not symmetric. There are two slightly different symmetrization methods that circumvent this problem. The more popular one, which we denote the ‘main’ method, uses an estimator that replaces the simple time-lagged averages above by averages over all pairs (**x**_*t*_, **x**_*t*+*τ*_) and (**x**_*t*+*τ*_, **x**_*t*_), following e.g. Noé [28] and the popular software package PyEMMA [29]. As a result, on the left hand side of equation (1) **C**(*τ*) is replaced with

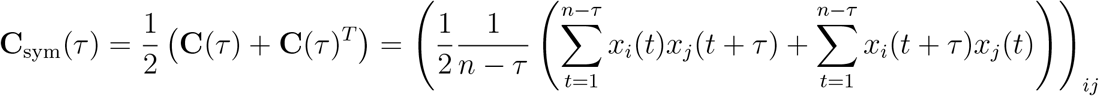

and on the right hand side **C**(0) with

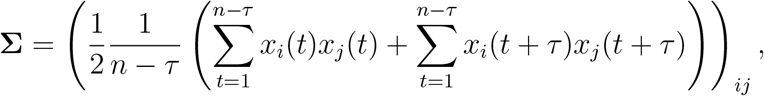

yielding a symmetrized version of equation (1) with real eigenvalues,

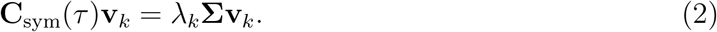

The second ‘alternative’ symmetrized version of equation (1) only differs on the right hand side, where **C**(0) is not replaced with **Σ**,

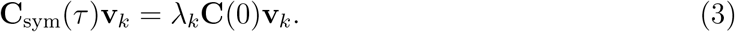

Our analysis is very similar for both versions, though with unexpectedly different results.

### 2.2 Theory

To render this symmetrized generalized eigenvalue problem more amenable to analysis, and following Ref. [30], we define a matrix formed from the trajectory

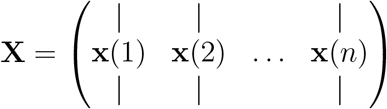

as well as a shorter time-lagged matrix

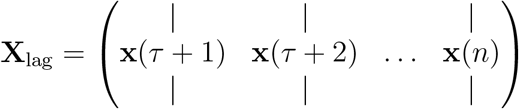

and one that is cut off at the end

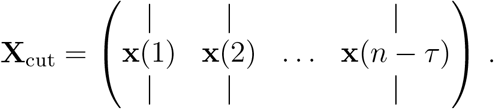

The latter two matrices serve to re-write the above left and right hand sides,

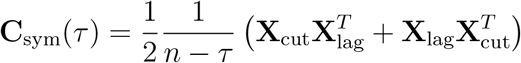

and

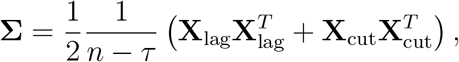

and, hence, also the symmetrized tICA-equation,

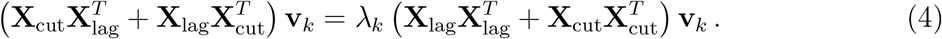

This defining equation (4) for tICA can be converted into a more convenient form using the

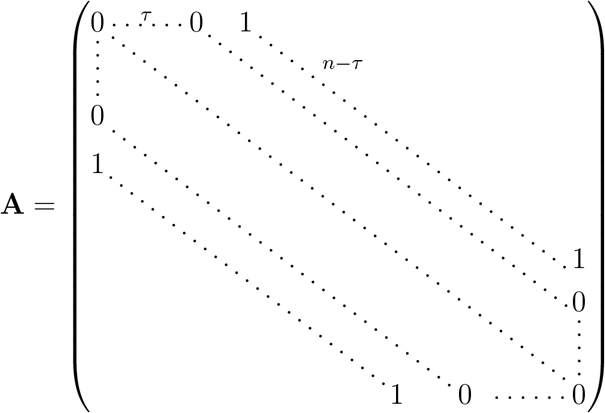

and

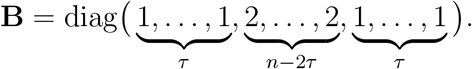

Noting that

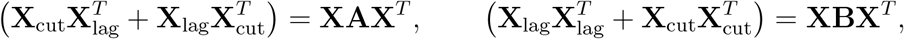

equation (4) reads

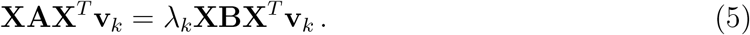

This can be transformed into a normal eigenvalue problem using the AMUSE-algorithm [31, 32] as follows. First diagonalize the right hand side by an orthogonal matrix **Q** and a diagonal matrix **Λ** such that

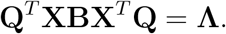

Substituting **v**_*k*_ = **Wu**_*k*_, with **W** = **QΛ**^−1/2^, and assuming all diagonal elements of **Λ** are nonzero, yields

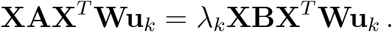

Note that this assumption is actually not necessarily true here, but since we are only interested in the nonzero eigenvalues and their eigenvectors the end results will still be correct. Since **W** is invertible, this equation is equivalent to

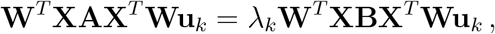

where the matrix on the right hand side turns out to be the unit matrix,

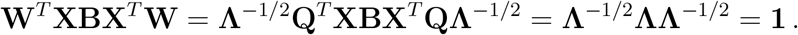

Hence equation (5) simplifies to

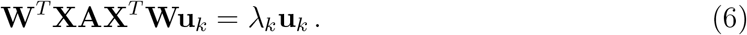

Now consider the following ‘swapped’ version [30]:

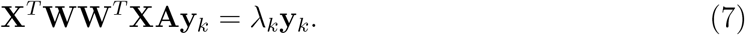

Notably, for each **y**_*k*_ satisfying equation (7) there exists a corresponding eigenvector that solves equation (6). Indeed, choosing **u**_*k*_ = **W**^*T*^**XAy**_*k*_ yields

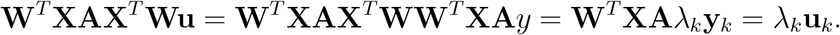

Finally, up to normalization, **y**_*k*_ is the projection of the trajectory onto the corresponding **v**_*k*_ = **Wu**_*k*_,

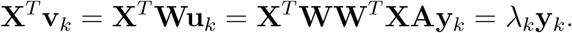

In other words, the tICA-projections of the trajectory are the eigenvectors (with non-zero eigenvalues) of the matrix **M** = **X**^*T*^**WW**^*T*^**XA**.

We will use this reformulation of the tICA defining equation to calculate the tICA-projections of random walks of given finite dimension and length.

### 2.3 Random Walks

For the numerical and semi-analytical evaluation of tICA components, random walk trajectories **x**(*t*) ∈ ℝ^*d*^ of dimension *d* were generated by carrying out *n* steps according to

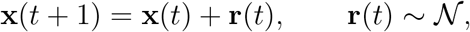

where 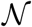 is a *d*-dimensional univariate normal distribution centered at 0. Each trajectory was centered to zero before further processing. We verified empirically that other fixed probability distributions with mean 0 and finite variance yield similar results.

### 2.4 Molecular Dynamics Simulation

For two proteins a 1 *μ*s molecular dynamics trajectory each was analyzed (Andreas Volkhardt, private communication). Both were generated using the GROMACS 4.5 software package [33] with the Amber ff99SB-ILDN force field [34] and the TIP4P-Ew water model [35]. The starting structures were taken from the PDB [36] entries 11AS [37] and 2F21 [38], respectively. Energy minimization was performed using steepest descent for 5 · 10^4^ steps. The hydrogen atoms were described by virtual sites. Each protein was placed within a triclinic water box using gmx-solvate, such that the smallest distance between protein surface and box boundary was larger than 1.5 nm. Natrium and chloride ions were added to neutralize the system, corresponding a physiological concentration of 150 mmol/l. Each system was first equilibrated for 0.5 ns in the NVT ensemble, and subsequently for 1.0 ns in the NPT ensemble at 1 atm pressure and temperature 300K, both using an integration time step of 2 fs. The velocity rescaling thermostat [39] and Parrinello-Rahman pressure coupling [40] were used with coupling coefficients of *τ* = 0.1 ps and *τ* = 1 ps, respectively. All bond lengths of the solute were constrained using LINCS with an expansion order of 6, and water geometry was constrained using the SETTLE algorithm. Electrostatic interactions were calculated using PME [41], with a real space cutoff of 10 Å and a fourier spacing of 1.2 Å. The integration time step was 4 fs, and the coordinates of the alpha carbons were saved every 10 ps, such that 10^5^ snapshots were available for each trajectory. Of these we discarded the first 10^4^ steps, leading to trajectories of length *n* = 9 · 10^4^.

## 3 Results and Discussion

To characterize the tICA components and projections of random walks, we will proceed in two steps. We will first analyse a special case, for which some analytical results can be obtained. Second, we will use the obtained insights to generalize this result to random walks of arbitrary length *n* and dimension *d* using a combined analytical/numerical approach. Subsequently, we will compare the obtained random walk projections to tICA analyses of biomolecular trajectories.

### 3.1 A Special Case

To gain first insight into the tICA components of a random walk, first consider the special case *d* = *n*, which allows for an almost fully analytical approach. In this case, all matrices in equation (7) are square and, assuming that **X** is invertible,

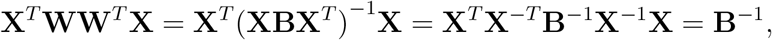

such that equation (7) becomes independent of **X**,

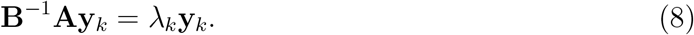

Note that the assumption that **X** is invertible is not strictly correct, as it has one zero-eigenvalue associated to the eigenvector given by **y**_0_ = (1, … , 1)^*T*^ . This is also an eigenvector of **B**^−1^**A**, but instead with eigenvalue 1. Therefore all the eigenvectors and all but one eigenvalue of equation (7) are identical to those of equation (8), and the analysis can proceed using equation (8).

In the limit of large *n*, and using the above definitions for **A** and **B**, the matrix **B**^−1^**A** approaches a circulant matrix with the property that each of its columns is a cyclic permutation of the preceding one. It differs from a circulant matrix only at the four ‘corners’ (of size *τ*) of the matrix, and for large *n* = *d* these ‘corners’ become small relative to the size of the matrix. More precisely, **B**^−1^**A** and the circulant matrix are asymptotically equivalent as in defined in Ref. [42].

Circulant matrices are diagonalized by the Fourier transform [43], yielding eigenvectors are

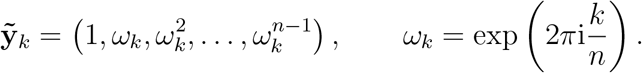

and eigenvalues

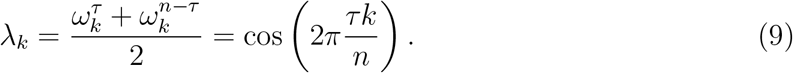

These eigenvectors are complex, but since *λ*_*k*_ = *λ*_*n*−*k*_ and 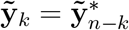, the real and imaginary part of 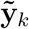 (cosine and sine) are real eigenvectors for the same eigenvalues. Depending on *τ* and *n*, many of these eigenvalues are equal, since they only depend on *τk* mod *n*.

This result implies that for large *n* = *d* the eigenvalues of **B**^−1^**A** approach those of the circulant matrix. More precisely, their eigenvalues asymptotically equally distributed [42]. In contrast, the eigenvectors are only preserved in limits or under small perturbations if the respective adjacent eigenvalues are well-separated from each other [44]. For the case at hand, however, this eigenvalue separation very quickly approaches zero for small *k* and large *n* (and for other *k* with | cos(2*πτk*/*n*)| ≈ 1). As a result, the eigenvectors of **B**^−1^**A** for small *k* (and other *k* as before) differ from those of the circulant matrix even in this limit. Rather, they need to be represented as approximate linear combinations of those eigenvectors of the circulant matrix with similar eigenvalues.

This subtlety contributes to the complexity of the problem as well as of the solution, and has so far prohibited us from proceeding further purely analytically both for finite *d* = *n* as well as for *d* = *n* → ∞. Nevertheless, the eigenvalue problem equation (8) provides a good starting point for a numerical approach. Still, the degeneracy discussed above needs to be taken properly into account, as the numerical eigenvectors are essentially arbitrarily chosen from the eigenspaces.

Inspecting the Fourier transforms of the numerical eigenvectors suggests that the eigenspaces of equation (8) for small *k* each contain an eigenvector that resembles a cosine function

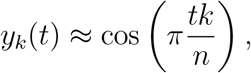

with increasing accuracy for increasing *n*.

Another effect of the poor separation of the eigenvalues is that the above results are very sensitive to small changes to the matrix in equation (8). E.g., using the alternative symmetrization method defined by equation (3), the analysis in Section 2.2 is unchanged, except that all diagonal entries of *B* become 2, and equation (8) reads

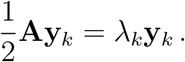

For *n* = *d* → ∞, the same circulant matrix is obtained, such that the eigenvalues, equation (9), are unchanged. The numerical solution however reveals that the first few eigenspaces instead contain eigenvectors given by

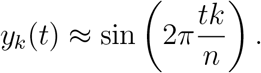

This result is indeed strikingly different, in that the cosine functions are replaced by sine functions with twice the frequency.

### 3.2 General Solution

Next, we will consider the general case, i.e., a random walk of length *n* in *d* < *n* dimensions. Unfortunately, we were unable to find analytical solutions similar to the above; however, the results of Section 2.2 permit an elegant way for a numerical approach by computing the expectation value of the matrix **M**. To this aim, **M** was computed for a sample of 20000 random walks of given fixed dimension *d* and number of time steps *n*, from which an average matrix ⟨**M**⟩ was computed. The eigenvectors of ⟨**M**⟩ served as the semi-analytical solution for the general case. We note that this does not necessarily produce the same results as averaging the individual tICA-projections directly. We have, however, tested that the eigenvectors of ⟨**M**⟩ are very similar to the averages of the tICA-projections. An exception to this is that averaging the tICA-projections can produce artefacts arising from to the fluctuating order of the eigenvectors, and these artefacts are not present in the eigenvectors of ⟨**M**⟩.

As an illustration, Figure 1 shows the first two resulting tICA-projections for random walks with *n* = 1000 and *d* = 50, revealing a strong dependence on the lag time *τ* . For short lag times *τ* , *y*_1_(*t*) ≈ cos(*πt*/*n*) and *y*_2_(*t*) ≈ cos(2*πt*/*n*). With increasing *τ* , this low-frequency cosines are gradually replaced by higher-frequency components, first in **y**_2_ (starting at about *τ* = 90) and for further increasing *τ* > 150 also in **y**_1_. From then on, the frequencies of both **y**_1_ and **y**_2_ slowly decrease, maintaining a *π* phase shift.

**Figure 1:**
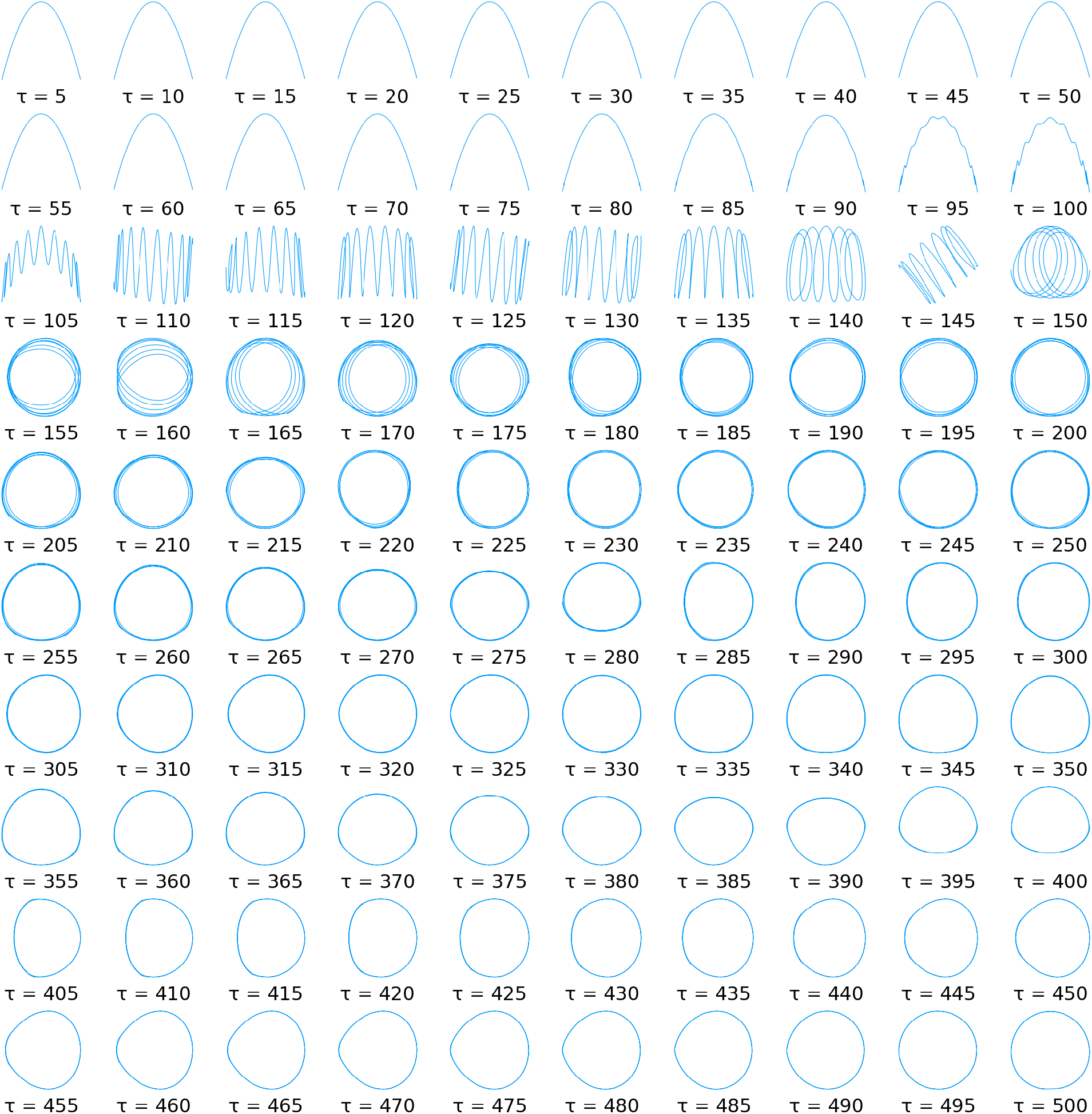
The first two ‘expected’ tICA-projections of random walks of dimension *d* = 50 with *n* = 1000 time steps for varying lag time *τ* , computed with the averaging method from Section 3.2 using a sample of 20000 random walks. For each *τ* , the first tICA-projection is shown on the x-axis and the second one on the y-axis.

In contrast to the special case considered above (Section 3.1), our numerical studies suggest that for large lag times the averaged projections do not approach exact cosines for large *n*. Rather, ‘cosine like’ functions appear, as can be seen for the high lag-times shown in Figure 1, where the circular shape that would be expected for exact cosines is noticeably distorted, even if *n* is further increased. In contrast, for short lag times, where the higher frequency components have not yet appeared (e.g. *τ* < 90 in Figure 1), the projections do seem to approach exact cosines with increasing *n*.

For the alternative symmetrization method, equation (3), the same method can be applied, and the obtained projections are shown in Figure 2. Indeed, comparing the two Figures, even more dramatic differences are seen as a result of this very small change. In particular, for short *τ* values, the cosine-like functions seem to be replaced by sine-like functions of twice the frequency, just like we have already seen for the special case *d* = *n*. Also, for increasing *τ* a much richer and complex behavior is seen. Finally, the onset of higher frequencies occurs for somewhat smaller *τ* values (at *τ* ≈ 100) compared to Figure 1 (at *τ* ≈ 110). This abrupt emergence of higher frequencies deserves closer inspection.

**Figure 2:**
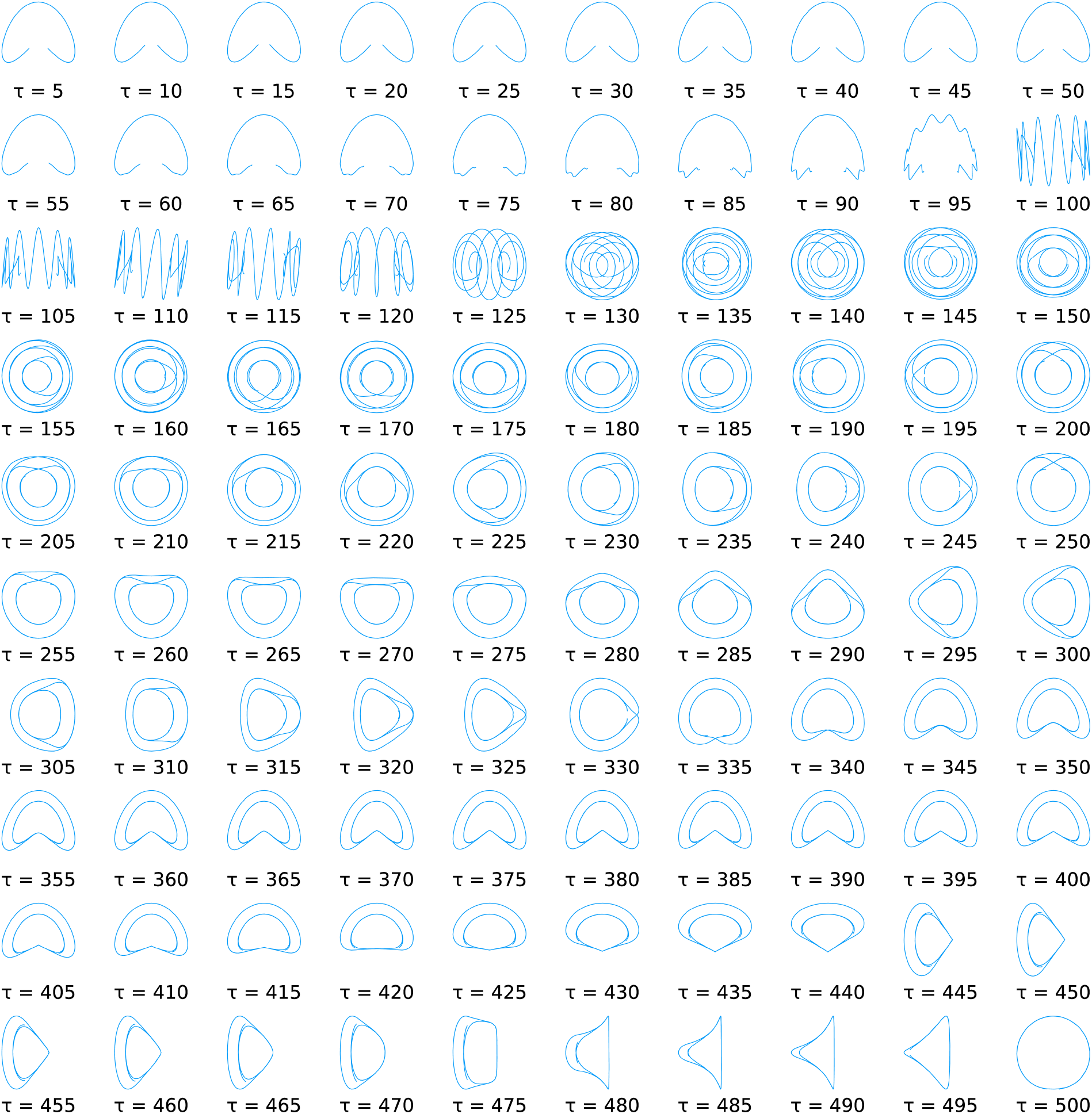
The first two ‘expected’ tICA-projections, for the alternative symmetrization method, of random walks of dimension *d* = 50 with *n* = 1000 time steps for varying lag time *τ* , computed with the averaging method from Section 3.2 using a sample of 20000 random walks. For each *τ* , the first tICA-projection is shown on the x-axis and the second one on the y-axis.

### 3.3 Abrupt Changes

To gain more insight into why these abrupt changes occur, Figure 3 (A) shows the eigenvalues of ⟨**M**⟩ as a function of *τ* for dimension *d* = 30, revealing a strikingly complex pattern. For small lag times *τ* all eigenvalues decrease with *τ* , with associated cosine-shaped eigenvectors of period lengths 2*n*, 2*n*/2, 2*n*/3, … , as annotated in the Figure. The decrease of these curves reflects the sampling of the cosine-shaped eigenvectors with increasing lag time *τ* and, hence, the respective autocorrelations also resemble cosine functions.

**Figure 3:**
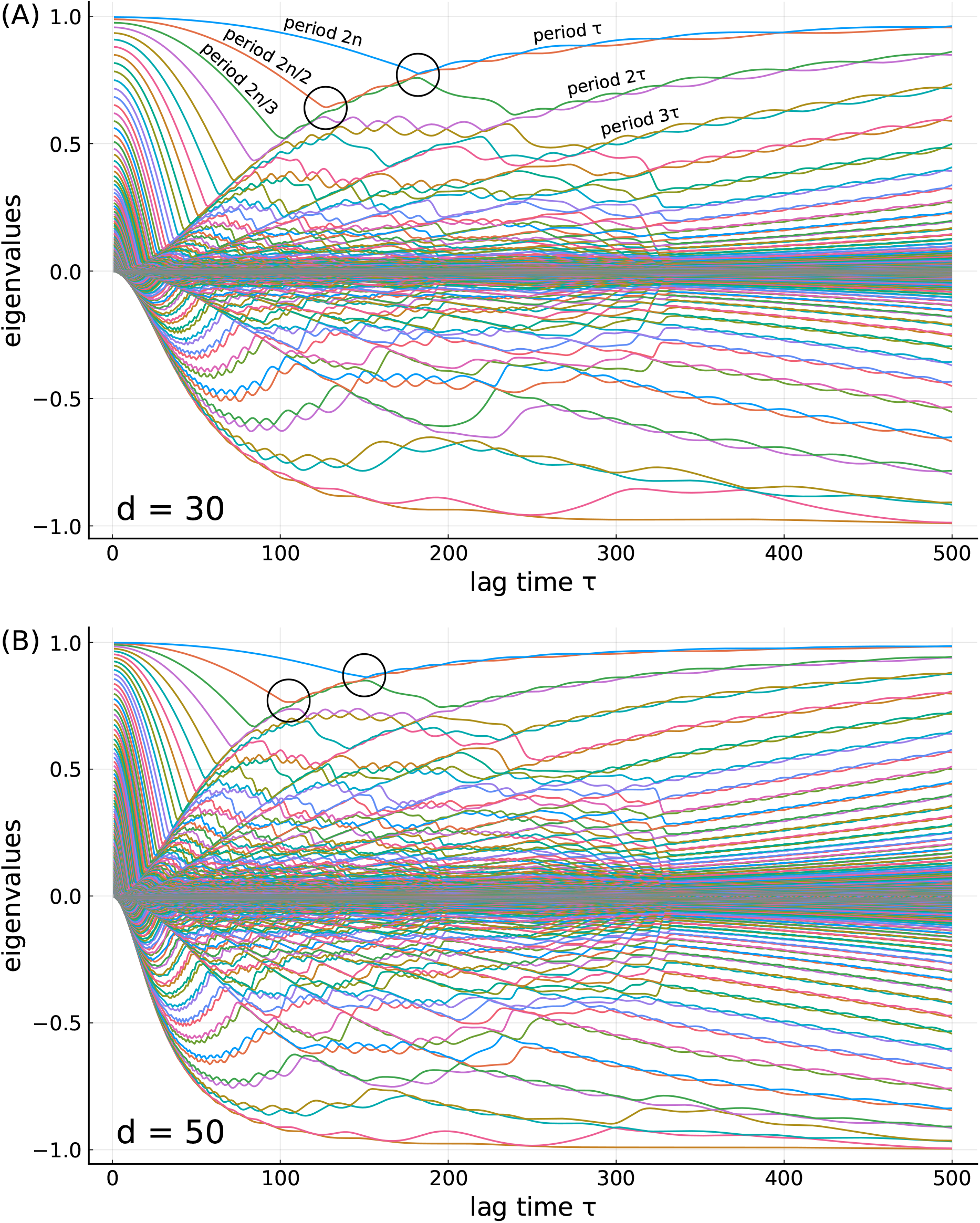
The eigenvalues of the averaged matrix ⟨**M**⟩ as a function of the lag time *τ* at (A) dimension *d* = 30 and (B) dimension *d* = 50. The two abrupt changes are indicated using black circles. The colors indicate the order of the eigenvalues.

Also visible are several curves that monotonically increase with *τ*, each starting at zero for small *τ*. These curves represent two eigenvalues each, with cosine-shaped and sine-shaped eigenvectors of period lengths *τ*, 2*τ*, 3*τ*, … , respectively, as also annotated in the Figure. Their increase is less obvious, as one might expect the autocorrelation of a *τ*-periodic function at lag time *τ* to be unity and, therefore, constant. Note, however, that the eigenvalue of ⟨**M**⟩ does not strictly represent this autocorrelation; rather, it represents the average of the autocorrelations of many instances of this eigenvector for each single random walk — each of which is not strictly periodic. For increasing period lengths, the eigenvectors approach cosines or sines, such that their average autocorrelation increases and so do the corresponding eigenvalues of ⟨**M**⟩.

At the intersections of these two sets of curves (black circles) the respective eigenvalues are degenerate and their order changes, which causes abrupt changes of the eigenvectors and, therefore, also of the projections onto these eigenvectors, the first two of which were discussed above.

For larger dimensions *d*, e.g., for *d* = 50 as shown in Figure 3 (B), one would expect that the tICA-projections resemble cosine or sine functions increasingly closely, also also at increasingly higher frequencies. As a result, the eigenvalues corresponding to the eigenvectors with period lengths *τ*, 2*τ*, 3*τ*, … should increase with *d* at any given lag time *τ* , whereas the decreasing eigenvalue curves on the left side should remain unchanged. Therefore, the respective intersections should occur at smaller lag times *τ* . Comparison of the black circles in the two panels of Figure 3 shows that this is indeed the case. To illustrate this effect, Figure 4 shows the first two tICA-projections of random walks with dimensions ranging from 50 (top row) to 500 (bottom row) for increasing *τ*.

**Figure 4:**
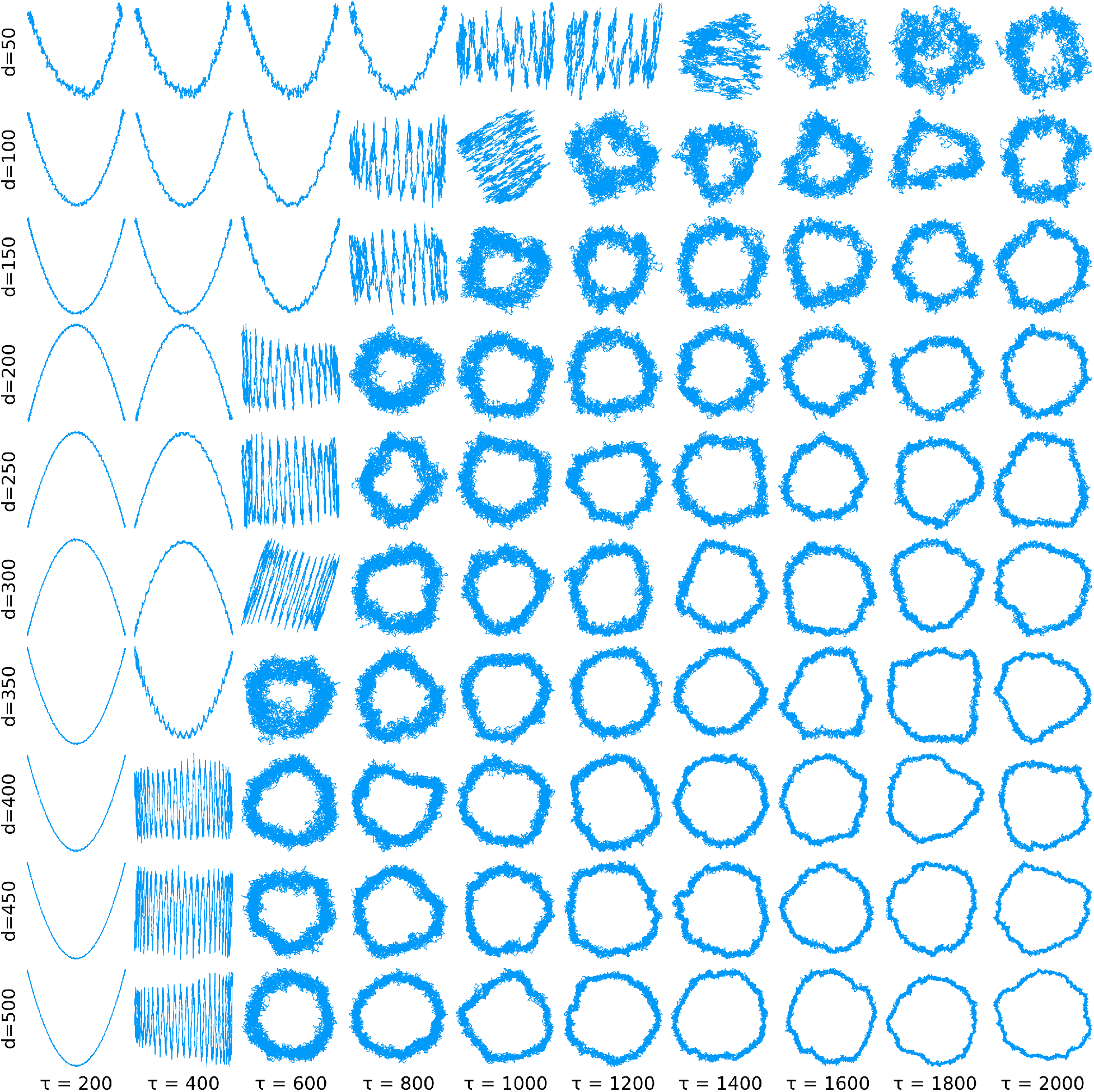
The first two tICA-projections of random walks with varying dimensions *d*, each with *n* = 10000. The lag times of the abrupt changes decrease with increasing dimension.

To quantify this behaviour, we generated a large number of random walks and determined the lag times *τ* at which the abrupt changes occur. Figure 5 shows the first and second of these critical lag times as a function of dimension *d* and for *n* ranging from 1000 to 5000 (colors). To enable direct comparison, the lag times *τ* have been normalised by *n*. As can be seen, for *d* between ca. 150 and *n/*2 both the first (upper curves) and second (lower curves) approximate power laws *n/τ* ∝ *d*^*b*^, as indicated by the respective fits (solid lines, the colors correspond to the values of *n*). For each fit, only dimensions *d* within the above range have been used.

**Figure 5:**
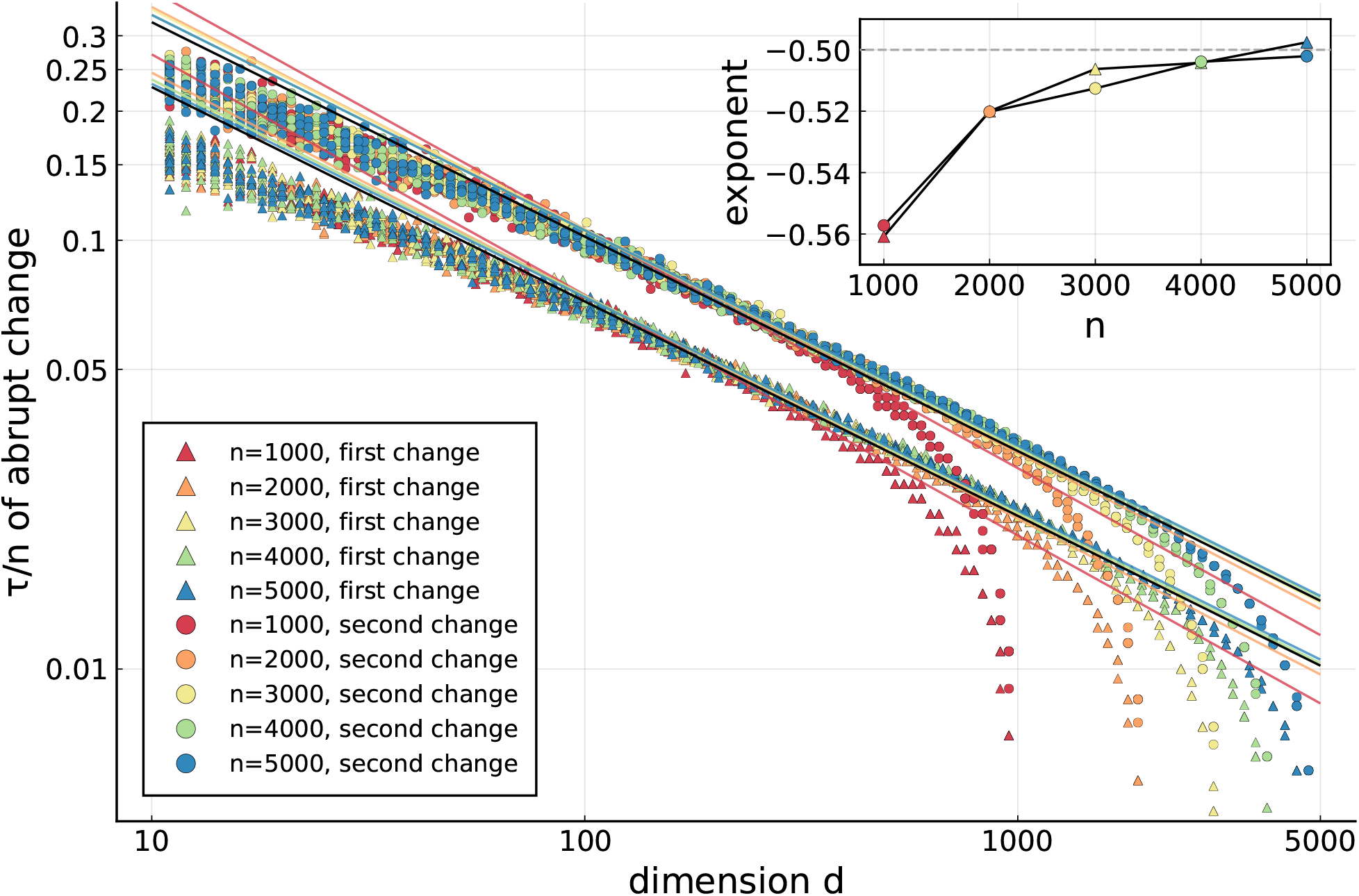
The lag time at which the abrupt changes occur in dependence of the dimension for various *n*. Each dot represents an independently generated random walk. Also shown are the power law fits *n/τ* = *a* · *d*^*b*^ (colored lines), their exponents (inset), and the lines corresponding to *b* = −0.5 (black lines).

The inset of Figure 5 shows the power law exponents *b* for varying *n* and for the first and second abrupt change, both of which apparently approach *b* = −1/2 for large *n* (also represented by the black lines in the main Figure). Although we were unable to find a rigorous proof, this finding suggests that in the limit of large *n* and *d*, with *d* markedly smaller than *n*, the first few lag times at which abrupt changes occur scale as 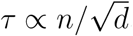.

### 3.4 Comparison of Random Walks and MD-trajectories

We next compared the tICA-projections of random walks with those of molecular dynamics trajectories of proteins in solution. To that end, we used two MD-trajectories of length 1 *μ*s each (generated as described in Section 2.4), one of a comparatively large protein (PDB 11AS, 330 amino acids) [37] and one of a smaller protein (PDB 2F21, 162 amino acids) [38].

As can be seen in Figure 6, the tICA-projections of the larger protein (top group) are indeed spectacularly similar to those of a random walk (bottom group). Even the strong dependence on the lag time is very similar, as are the abrupt changes discussed above.

**Figure 6:**
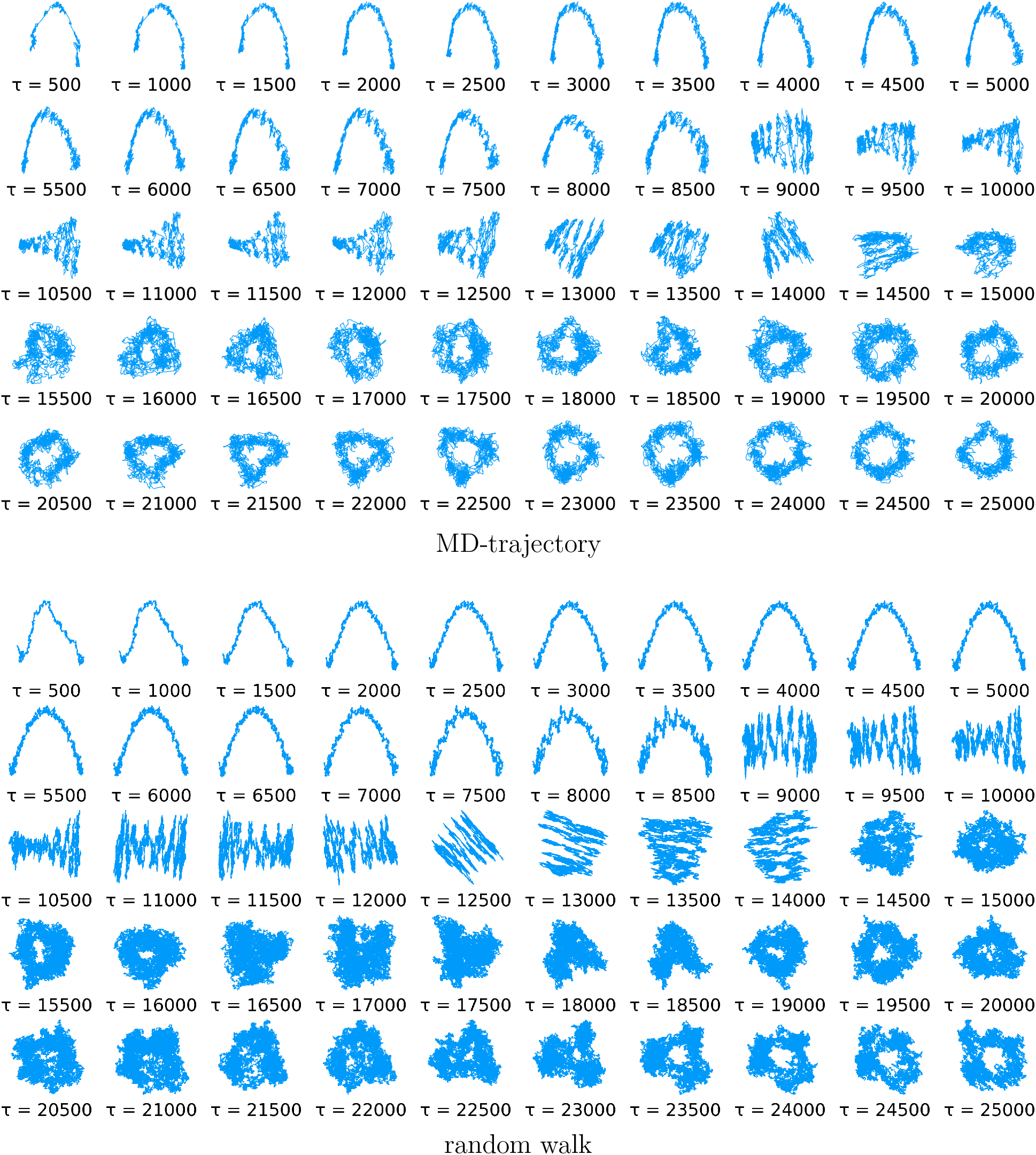
The first two tICA-projections of an MD-trajectory of PDB-entry 11AS (upper group) and those of a 40-dimensional random walk (lower group) for varying lag time *τ* . In this plot those of the MD-trajectory are smoothened using a moving average to improve readability.

Note that this striking similarity was obtained for a particular choice of *d* = 40 for the random walk; other dimensionalities yield less similar projections. Intriguingly, this finding thus suggests a new method of estimating an ‘effective’ dimensionality of MD trajectories.

It is also worth noting that both the MD-trajectory and the random walk projections show apparent ‘clusters’, e.g. for *τ* = 500 and *τ* = 8000, which also look quite similar. The fact that such clusters are also seen for the random walk strongly suggests that these are mostly stochastic artefacts and do not point to minima of the underlying free energy landscape.

Closer inspection of the random walk projections offers an additional possible explanation for some of the clusters, which may also apply to the MD trajectory projections. Focusing, e.g., at the averaged tICA-projections in Figure 1 immediately before the first abrupt change, one can see that the projection becomes overlayed with a cosine of higher frequency. Particularly at the ends of the curves, and in the presence of noise typical for single trajectories, this high frequency component can also produce apparent ‘clusters’.

In contrast, for the smaller protein (Figure 7) no similarity to the tICA-projections of random walks is observed. In fact, the tICA-projections of the trajectory of the smaller protein show no resemblance to a cosine-like function at all. In light of the above analysis, this finding suggests that this trajectory is sufficiently long to explore one or several minima of the underlying free energy landscape, thereby deviating from a random walk. Further, one may infer that the three clusters seen in the Figure actually point to conformational substates and, hence can serve as proper Markov states.

**Figure 7:**
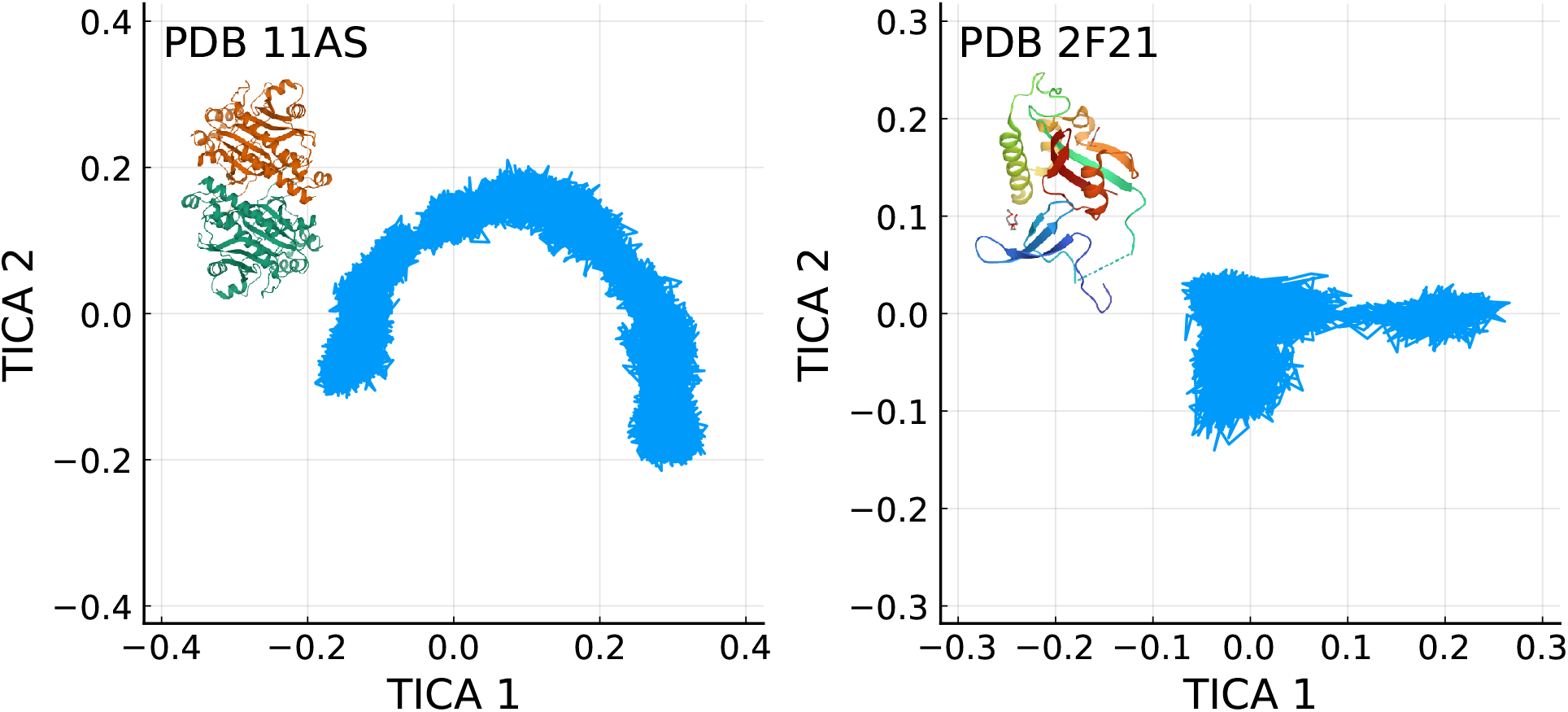
The first two tICA-projections of trajectories of the PDB-entries 11AS (on the left) and 2F21 (on the right). The larger protein (11AS) produces a cosine-like shape while the smaller one does not.

It is an intriguing question whether or not, for given trajectory length, larger or more flexible proteins tend to more closely resemble random walks.

## 4 Conclusions

Here we have analysed projections of random walks on tICA subspaces and subsequently compared those to tICA-projections of molecular dynamics trajectories of proteins. Our combined analytical and numerical study revealed a staggering complexity of the random walk tICA-projections, which showed a much richer mathematical structure than projections of random walks on principal components (PCA) [21, 22].

We attribute this complexity primarily to the fact that, in contrast to PCA, tICA components encode time information of the trajectory and, therefore, extract and process significantly more information. Mathematically, the complex behavior originates from the non-continuous switch of the order of eigenvalues for increasing lag time *τ* , when passing through points of eigenvalue degeneracy. At these points, the associated eigenvectors change abruptly, and so do the corresponding projections of both random walks and molecular dynamics simulations. We also find that tICA can be very sensitive to very small changes in the definitions of the involved matrices. In particular, the projections of random walks are very different for the two discussed symmetrization methods.

A particularly striking example is the first abrupt change of the projections onto the two largest eigenvalues. Here, a closer inspection revealed an approximate square root relationship between the lag times at which this occurs and the dimensionality of the random walk. A similar square root law is already known for PCA: Approximately the first 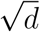 principal components of random walks resemble cosines [21].

Comparison of tICA-projections of random walks with those of a large protein (PDB 11AS) revealed striking similarities. This remarkable finding suggests that not only the ensemble properties of the finite protein trajectory resemble those of a random walk, as has been shown earlier via PCA [21], but also the time correlations of the underlying protein dynamics. Here, the appearance of cosine-like functions in the projections onto the tICA-vectors associated with the longest correlation times clearly points to a non-converged trajectory. For the comparatively small lag times typically used, the tICA-projections of random walks almost exactly resemble cosine functions, such that the cosine-content [22] of the tICA-projections should serve as a good quantifier of this.

In contrast, no resemblance to a random walk was seen for the second, smaller protein studied here, indicating that the projection reflects actual features of the underlying conformational dynamics of the protein.

The example in Figure 6 also illustrates the risk of over-interpreting apparent ‘clusters’ seen in the tICA-projections as actual conformational substates [4, 16], which are defined as local minima of the protein free energy landscape that are sufficiently deep for the system to stay there for a certain amount of time [16]. Clearly, it is tempting to also see ‘clusters’ in the random walk projections, which, however, by the definition of the random walk as a diffusion on a flat energy landscape, cannot represent conformational substates. This finding raises concerns for using automated clustering algorithms to identify, e.g., folding intermediates or to characterize conformational motions from tICA-projections [45].

Because the additional parameter of a varying lag time provides a much richer structure and many instead of only one projection (as is the case for PCA), tICA resemblance to a random walk offers a much more sensitive tool to detect lack of convergence in MD trajectories of large biomolecules. Further, by adjusting the dimension of the random walk such as to maximise the similarity to a given MD trajectory, one can estimate the effective dimensionality of the underlying dynamics. The latter idea, as well as precisely how this ‘effective dimensionality’ can be defined, clearly deserves further exploration.

## 5 Acknowledgements

We thank Nicolai Kozlowski, Malte Schäffner and Andreas Volkhardt for very helpful discussions; and Andreas Volkhardt for providing the MD-trajectories for our analysis. This work was supported by the German Ministry of Education and Research, BMBF project 05K20EGA and the German Science Foundation, grant SFB 1456.

This analysis has been implemented using the Julia programming language [46].

